# Simultaneous circulation and coinfection of Dengue clades and serotypes in Southern Brazil

**DOI:** 10.1101/2024.02.24.581892

**Authors:** Thales Bermann, Ludmila Fiorenzano Baethgen, Tatiana Schäffer Gregianini, Fernanda Godinho, Regina Bones Barcellos, Amanda Pellenz Ruivo, Milena Bauerman, Taina Machado Selayaran, Franciellen Machado dos Santos, Julio Augusto Schoerer, Érica Bortoli Möllmann, Fernanda Letícia Martiny, Valeska Lizzi Lagranha, Luiz Felipe Valter de Oliveira, Ana Gorini da Veiga, Andreia Carina Turchetto Zolet, Gabriel Luz Wallau, Richard Steiner Salvato

## Abstract

Dengue is an arthropod-borne virus with worldwide urban transmission mediated by anthropophilic vectors of the genus *Aedes*. Currently, there is a substantial genotype diversity that belongs to at least four known serotypes. Tropical regions, where several serotypes cocirculate in the population, are hyperendemic, driving multiple outbreaks with variable disease manifestations. Notably, the Dengue virus burden is increasing in subtropical and temperate regions of the globe due to climatic changes associated with new outbreaks in naive and previously unaffected human populations. The introduction and endemic transmission of different clades and serotypes in these regions is highly concerning due to the known and still unknown consequences of Dengue infection on populations with diverse genetic backgrounds. Here, we characterized the coinfection of different Dengue clades and serotypes in Rio Grande do Sul, Brazil’s southernmost and subtropical state, by combining serotype-specific RT-PCR and deep sequencing. We found five coinfected patients in four municipalities where these clades and serotypes circulated simultaneously. Our data suggests that coinfection events account for a minority of total cases, but these findings may reflect the early transmission phases of some Dengue clades. Hence, close monitoring is warranted to evaluate coinfection rates and associated clinical consequences as clade prevalence changes.

## INTRODUCTION

Arthropod-borne viruses (arboviruses) pose a significant global threat, but due to the link with arthropod vectors, their incidence, and burden are particularly high in tropical regions of the globe (Marklewitz and Junglen, 2019). In the last years, the global arbovirus burden has increased substantially associated with global travel (spread of viruses originally restricted to some regions of the globe and spread into naive populations) and to the continuous expansion of the vectors to subtropical and temperate regions due to climate change (Mayer et al., 2017). Transmission to humans is primarily through blood-sucking insect vectors, predominantly mosquitoes and ticks. Brazil has witnessed numerous arbovirus outbreaks in recent years driven by Zika and Chikungunya virus (ZIKV and CHIKV) arrival, and country-wide spread on top of the decade-long persistence of dengue in the country (Brito et al., 2021; Gregianini et al., 2017; Kading et al., 2020). However, new arboviruses emerged and contributed to the overall arbovirosis burden, the Dengue virus (DENV) continues to be the most prevalent and geographically dispersed arbovirus in Brazil (Da Silva Neto et al., 2022). This scenario is due to the vast available suitable environments for mosquito breeding in urban settings and the large human populations living in poor sanitary conditions exposed to mosquito-borne pathogens over the past four decades. Moreover, the lack and limited availability of effective mosquito control methods, medicines, and vaccines also contribute to the largely unmitigated DENV outbreaks in the country (Villabona-Arenas et al., 2014).

DENV has been circulating in Brazil since the 1800s, but surveillance data has shown that successive large-scale outbreaks have become established in the country after a more recent DENV reintroduction around the 1980s (Barreto and Teixeira, 2008; Siqueira et al., 2005). Currently, Brazil is the most heavily affected country by dengue in the Americas, with over 3 million suspected cases investigated and more than 1.4 million confirmed cases in 2023. In this same period, dengue was responsible for more than 1,100 deaths in Brazil (https://datasus.saude.gov.br). Three states of the South region, historically less affected region of the country, concentrated a substantial proportion of these cases (Codeco et al., 2022). Rio Grande do Sul (RS), the subtropical southernmost state of Brazil, has experienced a significant surge in dengue cases over the past five years. In 2022, it recorded its highest disease incidence in the recorded historical series: the number of suspected cases reached 95,735, and confirmed cases totaled 67,322, yielding an incidence of 589.4 confirmed cases per 100,000 inhabitants. In 2023, a total of 73,285 suspected dengue cases were reported, of which 38,240 were confirmed, yielding an incidence of 341.4 confirmed cases per 100,000 inhabitants. This data confirmed a higher lethality rate, with fifty-four deaths recorded, compared to sixty-six deaths in 2022 (https://dengue.saude.rs.gov.br). Previous reports on Dengue in Rio Grande do Sul until 2022 revealed the presence of four serotypes circulating in different years (Tumioto et al., 2014) associated with many travel-associated introductions. Still, information in the following years is scarce and patchy and does not cover the spread of the disease at the state level (Gularte et al., 2023).

Molecular DENV surveillance in Rio Grande do Sul on human samples has been underway since 2011 using different diagnostic protocols (Gregianini et al., 2018; Tumioto et al., 2014), but since 2016, arbovirus surveillance has been strengthened in the whole country as a response to the Zika and Chikungunya epidemics (Gregianini et al., 2017). Currently, ELISA-based and RT-PCR assays are being routinely used, and more recently, genomic surveillance has been systematically implemented. These methodologies provide different levels of information ranging from past and present viral infections to detailed genetic information required to access the most prevalent and fixed genome-wide mutations as well as fine-grained intra-host nucleotide changes generated during a single patient infection (Ko et al., 2018; Pollett et al., 2020, 2018). Intra-host variants are particularly interesting since they can reveal de novo mutations emerging and being maintained at low frequency and coinfection events of related viral clades that may change clinical manifestations (Dezordi et al., 2022b; Machado et al., 2019). Arbovirus coinfection has been reported in several places around the globe including different virus species (DENV, CHIKV and ZIKV) and DENV serotypes (Brito et al., 2017; Carrillo-Hernández et al., 2018; Furuya-Kanamori et al., 2016; Machado et al., 2019; Marinho et al., 2017; Rodriguez-Morales et al., 2016; Thai et al., 2012; Villamil-Gómez et al., 2016). However, it is still unclear how frequent coinfection is in different settings and if there are any implications in terms of disease severity.

In this study, we used RT-PCR, deep-sequencing, and intrahost variant analysis along with phylogenetic reconstruction to detect potential coinfection events among DENV in samples obtained from diagnosed patients in the Southernmost State of Brazil. Our results suggest that coinfection is rare, but the changing prevalence of cocirculating clades may change this scenario. Moreover, close monitoring is warranted to evaluate and characterize the clinical manifestation (i.e., antibody-dependent enhancement) of a higher number of coinfected patients in endemic settings with high transmission of multiple DENV clades, genotypes, and serotypes.

## METHODS

### Patient samples and DENV detection

This study included 8,951 serum samples received by the Rio Grande do Sul Central Laboratory of Public Health (LACEN□RS) from 2015 to 2023 for DENV detection and serotyping. Ribonucleic acid (RNA) was isolated from samples using automated extraction on Extracta 96 equipment (Loccus), using the Fast Kit – Viral DNA and RNA (MVXA-P096 FAST) (Loccus, São Paulo, Brazil) according to manufacturer’s instructions.

DENV detection and serotyping were performed through reverse transcription-quantitative polymerase chain reaction (RT-qPCR) using the Kit Molecular ZDC (Bio-Manguinhos, Brazil) according to the manufacturer’s instructions. Samples showing amplification with RT-qPCR cycle threshold (Ct) ≤38 were considered DENV-positive; positive and negative controls were used in each analysis. The multiple detection of the four known DENV serotypes (DENV-1 to 4) was performed in a CFX Opus Real-Time PCR System (Bio-Rad, Hercules, CA, USA).

### Genome sequencing, assembly, and minor variant analysis

Extracted RNA from 667 samples showing amplification with RT-qPCR Ct ≤28 was used for genomic sequencing. The genomic libraries were prepared using an adaptation of the Illumina COVIDseq protocol (https://www.illumina.com/products/by-type/ivd-products/covidseq.html), replacing the primers by previously published DENV primers sets (Brito et al., 2021; Gräf et al., 2023). Sequencing was performed using Illumina Miseq (Illumina Inc.). A reference-based genome assembly was conducted using the *ViralFlow 1*.*0* pipeline (Dezordi et al., 2022a), using the closest and most well-annotated DENV-1 (GenBank: GU131863.1) and DENV-2 (GenBank: MW577818.1) reference genomes.

The intrahost variant analysis was also performed using the ViralFlow pipeline, as previously described (Dezordi et al., 2022b). In brief, after consensus generation, the *bam-readcount* tool extracts the proportion of each base (A, C, T, G) at every position from the bam file. Subsequently, an in-house Python script (intrahost.py) is employed to identify intrahost single nucleotide variant (iSNV) sites based on specific criteria. The defined rules for iSNV identification include requiring the minor variant (MinV) to constitute at least 5% of the total position depth. Additionally, it should be present in both sense and antisense reads, with each contributing at least 5% to the overall variant frequency and a minimum depth of 100 reads. Following identifying iSNV sites, two consensus genomes are generated as output. The major variant (MajV) is derived from the nucleotide in the major allele frequency at each genomic position. On the other hand, the MinV is constructed based on the nucleotide found within the minor allele frequency at the same positions.

### Genotyping and phylogenetic analysis

Our samples were analyzed using the *Genome Detective Virus Tool Version 2*.*72* (https://www.genomedetective.com) to associate genotypes belonging to each sample. To characterize the transmission chains of the DENV clades, we downloaded all near complete DENV-1 and DENV-2 genome sequences with coverage breadth > 70% from NCBI (https://www.ncbi.nlm.nih.gov/), resulting in a data set of 3.475 DENV-1 and 3.090 DENV-2 samples. Genomes were aligned with sequences generated in this study using *MAFFT* (Katoh et al., 2019)and manually curated to remove artifacts using *AliView* (Larsson, 2014). Maximum likelihood (ML) phylogenetic trees were estimated using *IQ-TREE2* (Minh et al., 2020) under the General Time Reversible (GTR) nucleotide substitution model, which was inferred as the best-fit model by the *ModelFinder* application implemented in *IQ-TREE2* (Kalyaanamoorthy et al., 2017). Statistical support for tree nodes and branches was estimated using aLRT and ultrafast bootstrap approximation approaches with 1000 replicates. To understand the phylogenetic placement of MajV and MinV genomes, we added both genomes to our datasets to reconstruct an ML tree, including the sequences collected from GenBank.

## RESULTS AND DISCUSSION

### Study sampling

In the studied period (2015-2023) Rio Grande do Sul state registered 1,023,366 notified Dengue cases, totalizing 177,009 confirmed cases distributed in the following years: 2015 (1,299 cases), 2016 (2,442 cases), 2017 (24 cases), 2018 (28 cases), 2019 (1,346 cases), 2020 (3,638 cases), 2021 (10,601 cases), 2022 (67,326 cases) and 2023 (38,357 cases).

The Rio Grande do Sul Public Health Laboratory identified 3,984 DENV-positive samples using the RT-qPCR method over the nine years studied. The samples were sourced from individuals with suspected acute febrile illness from 2015 to 2023, collected across 239 cities in the state, and exhibited amplification with RT-qPCR Ct of ≤38. We obtained raw reads of 627 samples diagnosed with DENV-1 infection and 40 from DENV-2 infection, totaling 667 samples investigated through whole genome deep sequencing. The average coverage breadth across the genome from all samples ranged from 72.23% to 99.84%, and an average coverage depth of 1817.46 (SD=908.59), ranging from 194.0 to 6,625.5.

### Serotype coinfection detection

During the study period, we also identified the coinfection of serotypes DENV-1 and DENV-2 identified by RT-qPCR in a sample from a 52-year-old female patient without risk factors. The patient lives in a northwest municipality of the Rio Grande do Sul state, where there was the simultaneous transmission of DENV-1 and DENV-2 serotypes. We sequenced both patient genomes to confirm serotype coinfection and performed phylogenetic reconstruction using maximum likelihood. The genome coverage breadth was 99.22% for DENV-1 and 91.57% for DENV-2. Additionally, the average coverage depth was 286.0 for DENV-1 and 770.0 for DENV-2. The results confirmed the coinfection by allocating each genome to its corresponding serotype in the tree, with maximum bootstrap support values for the two main branches representing the two serotypes, as shown in **Figure 1**. More data on samples Denv1_16248_2023 and Denv2_16248_2023 are presented in the following discussion and **Table 1**.

**Table 1.**
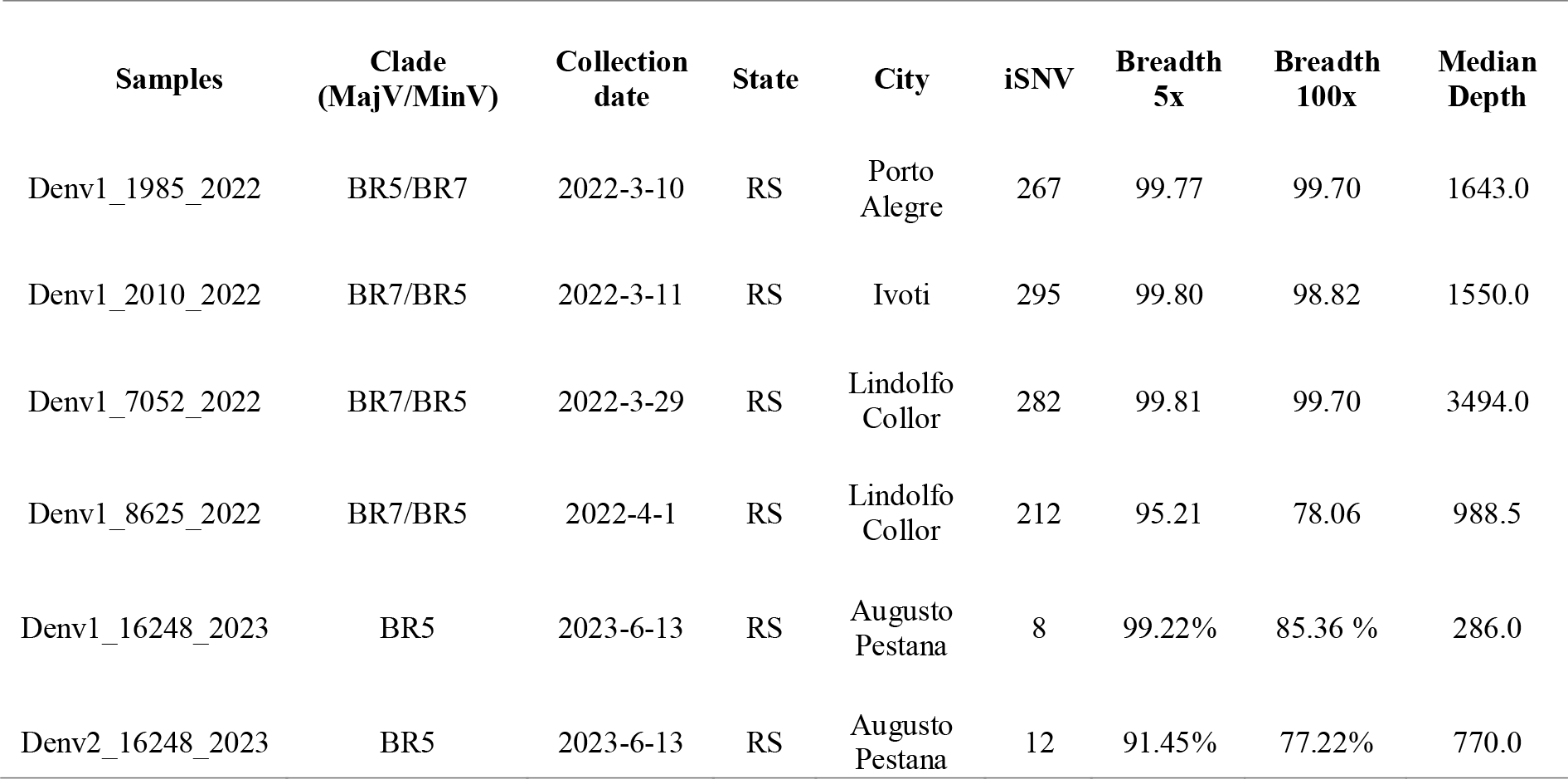
Summary of codetection samples.

| Samples | Clade<br>(MajV/MinV) | Collection<br>date | State | City | iSNV | Breadth<br>5x | Breadth<br>100x | Median<br>Depth |
| --- | --- | --- | --- | --- | --- | --- | --- | --- |
| Denv1_1985_2022 | BR5/BR7 | 2022-3-10 | RS | Porto Alegre | 267 | 99.77 | 99.70 | 1643.0 |
| Denv1_2010_2022 | BR7/BR5 | 2022-3-11 | RS | Ivoti | 295 | 99.80 | 98.82 | 1550.0 |
| Denv1_7052_2022 | BR7/BR5 | 2022-3-29 | RS | Lindolfo Collor | 282 | 99.81 | 99.70 | 3494.0 |
| Denv1_8625_2022 | BR7/BR5 | 2022-4-1 | RS | Lindolfo Collor | 212 | 95.21 | 78.06 | 988.5 |
| Denv1_16248_2023 | BR5 | 2023-6-13 | RS | Augusto Pestana | 8 | 99.22% | 85.36 % | 286.0 |
| Denv2_16248_2023 | BR5 | 2023-6-13 | RS | Augusto Pestana | 12 | 91.45% | 77.22% | 770.0 |

**Figure 1.**
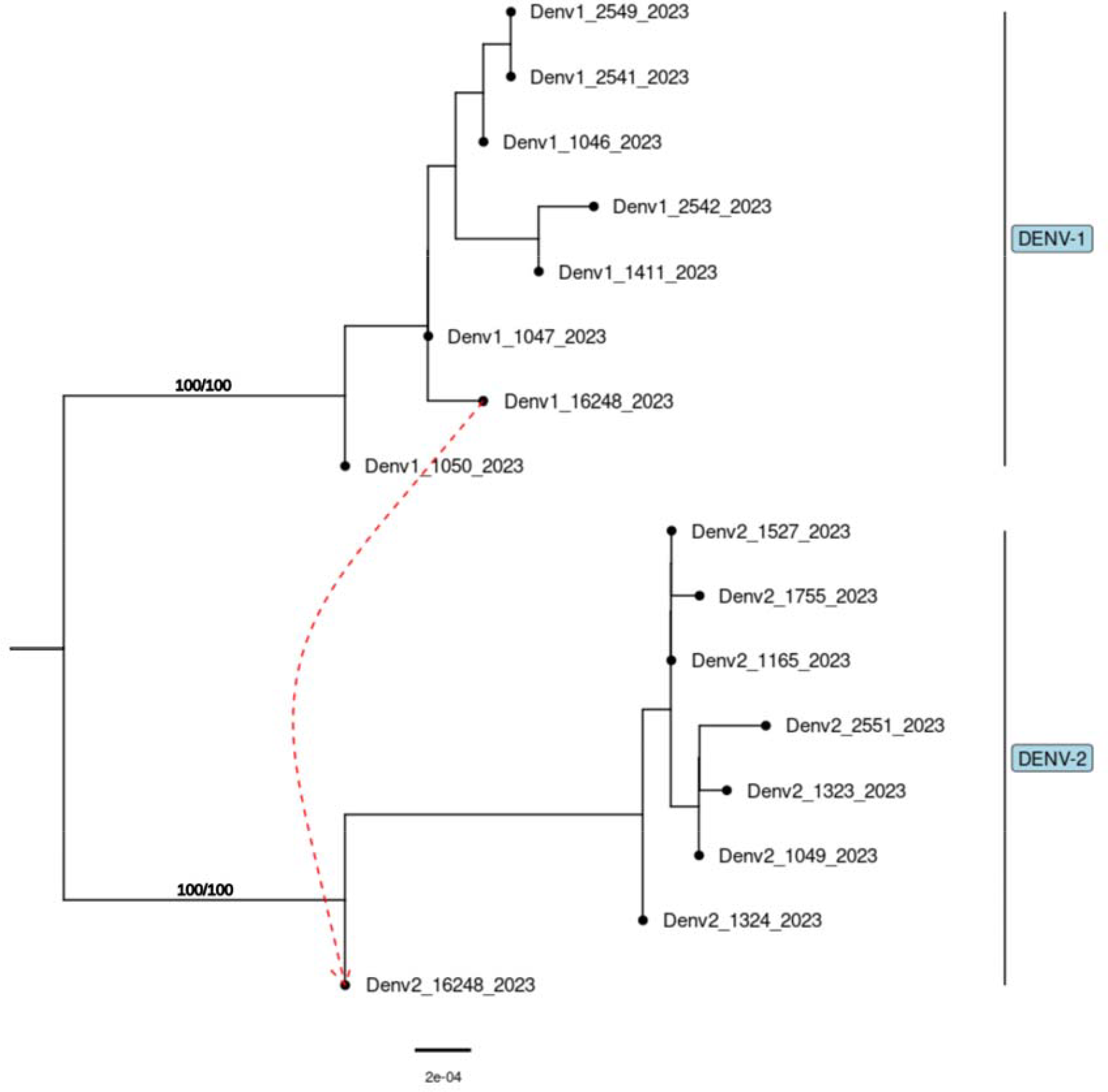
Maximum likelihood tree generated from a representative subset of the total dataset (total dataset tree for DENV1 and DENV2 are available in supplementary materials, **supplementary figure 1**). The red dashed line links the patient and both genomes between the two serotypes identified on the right side. Support values were estimated at 100/100 bootstraps in the two main branches.

### Minor variant analysis

Among the 667 genomes, 665 (99.70%) displayed at least one iSNV site, that is, at least one genomic position with more than 100 reads supporting a minimum of two alternative nucleotides. For DENV-1, an average of 20 iSNVs sites per sample was observed. For DENV-2, an average of 14 iSNVs was observed in the samples analyzed. Most samples (663) showed a range between 1 to 88 iSNVs. Interestingly, four DENV-1 samples showed a higher number of iSNVs between 212 and 295, suggesting possible coinfection, with a notable distance iSNV number gap between those and the remaining 663 samples. Notably, in the case of the serotype coinfection sample, both genomes Denv1_16248_2023 and Denv2_16248_2023 exhibited low iSNVs values (8 and 12, respectively), indicating a low intrahost diversity. The iSNVs values for all samples are available in **Supplementary Table 1**.

With few exceptions, most studies estimating DENV iSNV diversity have used cloning and Sanger sequencing (Descloux et al., 2009; Parameswaran et al., 2012; Wang et al., 2002). By comparing our data, it is suggested that the intrahost genetic variability of DENV is lower than previously estimated, as determined by Sanger sequencing, probably due to the more stringent parameters to validate low-frequency variants obtained through deep sequencing. For DENV-2, the intra-host variability found in this study showed a range of 2 to 43 iSNVs, reinforcing previous findings from deep sequencing, where a range of 10 to 44 iSNV was found for the complete DENV-2 genome (Romano et al., 2013). With the lack of comparable data from deep sequencing, we found a study evaluating intrahost variability for DENV-1 using cloning and Sanger sequencing, where they applied rigorous methods to validate low-frequency variants, obtaining an average of 15.61 iSNVs leading to a result consistent with the reports in this study for DENV-1 (Thai et al., 2012).

The unusually high number of iSNVs found in four DENV-1 samples may be derived from de novos SNPs generated during viral replication or due to simultaneous infections of clades having enough differentiating SNPs. Both phenomena would result in high iSNV, but using the MajV and MinV genomes generated, we sought to distinguish between these hypotheses through phylogenetic analysis of minor and major reconstructed genomes. Coinfection is expected to generate two substantially different genomes that confidently cluster within different clades, while de novo-generated minor and major reconstructed genomes are not. These four outlier samples and serotype coinfection data are described in **Table 1**. To assess if codetection could result from sample contamination, we assessed the negative controls from extraction to sequencing runs present in the runs of these four outlier samples. All the negative control assemblies revealed a coverage breadth and depth of 5% and 5x, respectively.

### Genotyping and phylogenetic analysis

MajV and MinV consensus sequences were generated for all samples bearing well-supported alternative nucleotides. Our analyses using *Genome Detective* showed that all 627 samples of DENV-1 belong to genotype V. Among the 40 samples of DENV-2, 33 were identified as genotype II - cosmopolitan, while seven were classified as genotype III - Southeast Asian/American. We found no difference between the MajV and MinV genomic sequences for these DENV2 genotypes.Phylogenetic analyses unveiled the presence of six distinct clades within our DENV-1 samples (BR3, BR4, BR5, BR6, BR7, and BR8), as shown in **Figure 2A**. Additionally, our analyses revealed the presence of two distinct clades within the DENV-2 samples (BR4 and BR5), as shown in **Figure 2B**. Among the six identified DENV-1 clades in our study, three (BR3, BR4, and BR5) were previously documented by Brito et al., 2021. Similarly, one of the two DENV-2 clades identified in our analysis (BR4) was previously characterized in the same study. Meanwhile, the DENV-2 (BR5) belonging to genotype II - cosmopolitan was reported in the state by Gräf et al., 2023.

**Figure 2:**
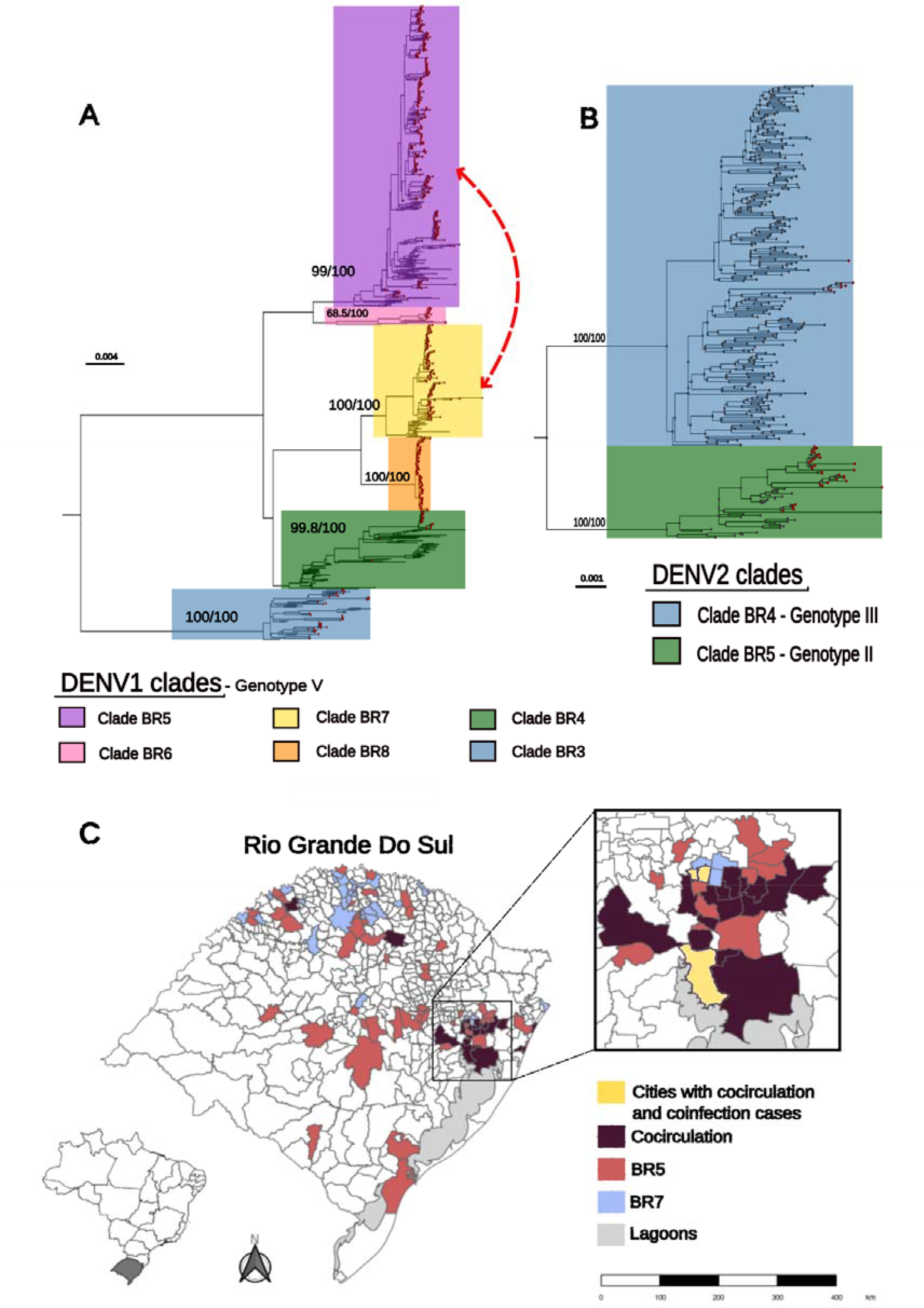
**A**. Maximum likelihood tree for DENV-1. The six clades identified in our study are highlighted, as shown in the legend below the tree. The red dashed line indicates the clades involved in coinfection cases; **B**. DENV-2 maximum likelihood tree. The two clades identified in our study are highlighted, as shown in the legend below the tree; Our samples are identified with a red dot. The bootstrap support values of the branches that lead to the clades are in figure; **C**. Map of the state Rio Grande do Sul, Brazil. Here, we show the dispersion of the BR5 and BR7 clades in our samples and the municipalities where they cocirculate. The zoom on the right side focuses on cocirculation in the capital’s metropolitan region, Porto Alegre, in addition to showing the three municipalities where we had the cases of coinfection reported in this study (they are also cocirculation zones), according to the legend.

Phylogenetic analysis revealed that MajV and MinV from the four outliers samples (Denv1_1985_2022, Denv1_2010_2022, Denv1_7052_2022 and Denv1_8625_2022) clustered with high branch support (using aLRT and ultrafast bootstrap approximation to assess branch supports) within different clades, confirming a coinfection by different clades for the four outlier samples. Coinfection occurred between two clades, BR5 and BR7. Denv1_2010_2022, Denv1_7052_2022, and Denv1_8625_2022, the MajV clustered within the BR7 clade, while their MinV genomic versions were clustered within the BR5 clade. The MajV genome of Denv1_1985_2022 clustered within the BR5 clade and MinV was placed within the BR7 clade (**Table 1**). The BR5 clade was described for the first time in 2021 by Brito et al. and is the most prevalent clade in our samples (51.38%). Its circulation in Rio Grande do Sul and Brazil was detected for the first time in 2015, being present and well dispersed in the state until the last sequenced samples of this study in 2023. The BR7 clade, also found in Peru in 2021, appears as an emerging clade in Rio Grande do Sul in 2022, present at an average frequency in our samples (19.61%).

The plausibility of the codetection events was further supported by the fact that the BR5 and BR7 clades identified by our study circulated simultaneously in the municipalities where the coinfection cases were found (**Figure 2C**). The four samples were sampled within a short time frame, with the first being collected on March 10, 2022, and the last on April 1^st^, 2022, with a distance of 21 days between them. All cases of clade coinfection were detected in 2022, where the two clades circulated in the metropolitan region of the state capital, Porto Alegre. The greatest geographical distances between the four samples are 60.9 kilometers from Porto Alegre to Lindolfo Collor. Notably, the occurrence of the four clade coinfection events coincided with a peak in observed cases in the region, emphasizing the plausibility of these detections. This context is further supported by the understanding that virus coinfection events are more likely to occur in endemic areas with cocirculation of different virus lineages (**Figure 3**).

**Figure 3.**
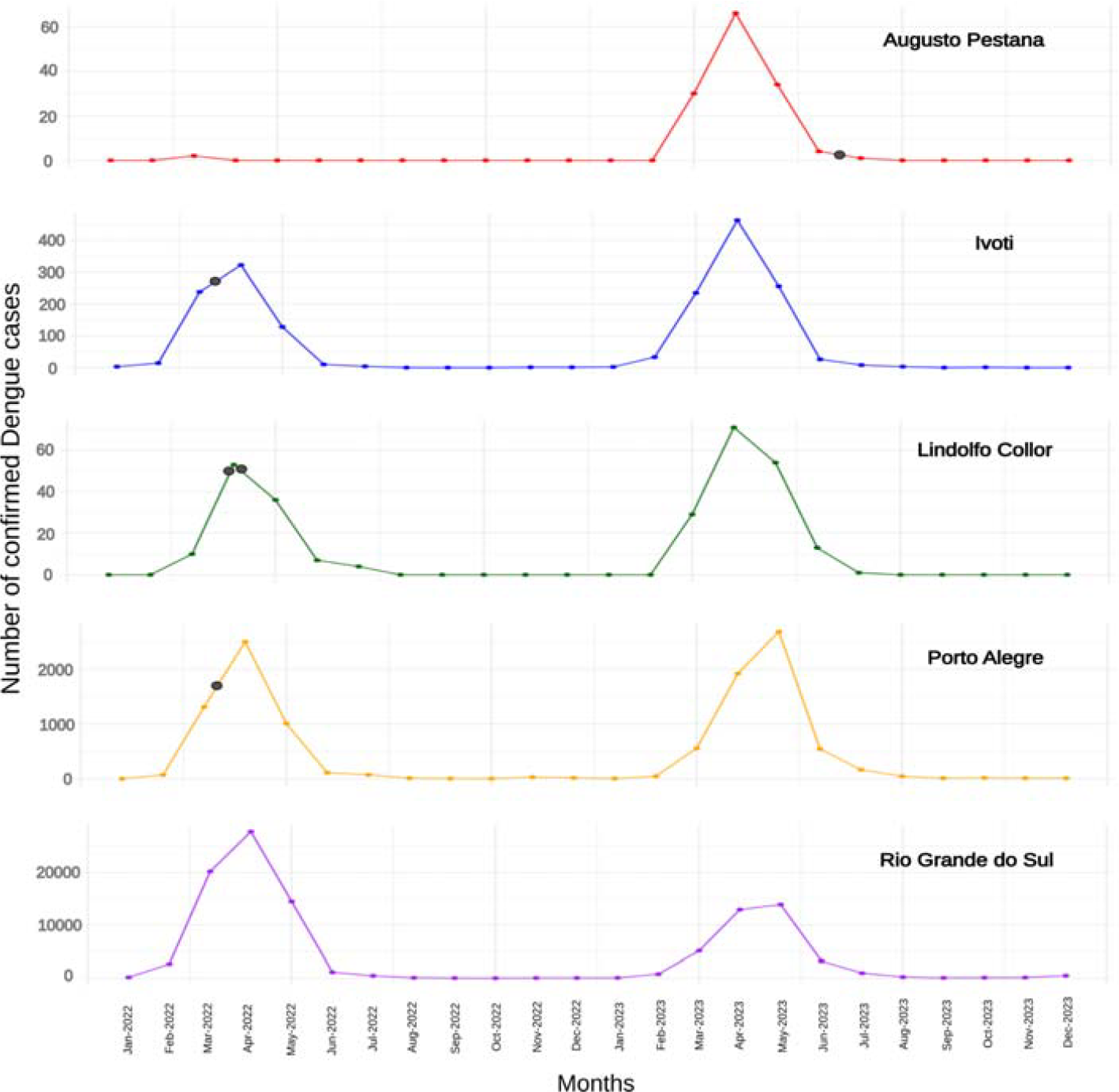
Plot with the number of confirmed dengue cases in cities and state-wide, where there were coinfection cases for 2022 and 2023. The gray dot represents the five coinfection events referenced temporally on the X axis.

Our current understanding of infection by different DENV clades and the occurrence of antibody-dependent enhancement (ADE) remains incomplete, with a lack of evidence for its occurrence and potential variation in the clinical outcome. Antibodies that developed during infection bind to DENV serotypes during subsequent infections. These antibodies activate the virus, forming a complex that binds to the Fcγ receptor (FcγR) on circulating monocytes. This interaction increases the ability of the virus to infect monocytes, leading to viral replication and an increased risk of severe dengue disease (Halstead, 2014; Shukla et al., 2020). Addressing these knowledge gaps is needed to develop effective strategies to prevent and treat dengue. The symptoms of the patient coinfected by serotypes DENV-1 and DENV-2 were fever, headache, myalgia, nausea, back pain, retroorbital pain, arthritis, and leukopenia. The dual infection seems not to have caused a severe dengue clinical presentation in this patient since no hospitalization was required and no hemorrhagic manifestation or other more severe alterations occurred. Similarly, all four identified cases of coinfection by distinct clades presented only mild symptoms and did not require hospitalization.

## CONCLUSION

Our in-depth analysis revealed at least five codetection events, corroborated by epidemiological data from geographically and temporally cocirculating clades in Rio Grande do Sul. Our results suggest that DENV codetection/coinfection events are rare, occurring at a rate of 0.75% in the studied population. However, this is undoubtedly a lower-bound estimate due to the limitation of detecting coinfection events with poorly divergent viral lineages. These findings have important implications for understanding the epidemiology of DENV in our region. The detection of coinfection between different clades shows the complexity of viral dynamics and may have significant repercussions on disease control. For example, coinfection by different clades can lead to greater genetic diversity of the virus, potentially influencing the host immune response and disease severity; further studies may help to clarify the interactions between multiple clade infections and the role of ADEs in shaping the clinical outcome of infection. As observed in our study, identifying coinfections by clades that circulate simultaneously in a given region emphasizes the need for continuous monitoring and genomic surveillance to better understand virus spread and predict possible outbreaks. Our results showed the importance of deep sequencing and refined phylogenetic analysis to detect and characterize coinfection events by different DENV clades, providing valuable information to guide disease control and prevention strategies. These findings also highlight the continued need for research to improve our understanding of DENV viral dynamics and their public health impacts.

## FUNDING

This work was supported by Fundação de Amparo à Pesquisa do Estado do Rio Grande do Sul (FAPERGS/FIOCRUZ 13/2022 – REDE SAÚDE-RS, grant process 23/2551-0000510-7 and FAPERGS 14/2022 - ARD/ARC, grant process 23/2551-0000852-1). Conselho Nacional de Desenvolvimento Científico e Tecnológico (CNPQ) and Fundação de Amparo à Pesquisa do Estado do Rio Grande do Sul has provided a fellowship to R.S.S (FAPERGS 14/2022 - ARD/ARC).

## CREDIT AUTHORSHIP CONTRIBUTION STATEMENT

**Thales Bermann:** Conceptualization, Investigation, Writing – review & editing. **Ludmila F. Baethgen:** Conceptualization, Investigation, Writing – review & editing. **Tatiana S. Gregianini:** Conceptualization, Investigation, Writing – review & editing. **Fernanda Godinho**: Methodology, Investigation, Writing - Review & Editing. **Regina B. Barcellos:** Methodology, Investigation. **Amanda P. Ruivo:** Methodology, Investigation. **Milena Bauerman**: Methodology, Investigation. **Taina M. Selayaran:** Methodology, Investigation, Software. **Franciellen M. Santos:** Methodology, Investigation. **Julio A. Schoerer:** Data Curation, Formal Analysis. **Érica B. Möllmann:** Data curation, Methodology. **Fernanda L. Martiny**: Data curation, Methodology. **Valeska L. Lagranha:** Data Curation. **Luiz F.V. Oliveira:** Funding acquisition, Methodology, Resources. **Ana G. Veiga:** Funding acquisition, Writing – review & editing. **Andreia C.T. Zolet:** Supervision, Writing – review & editing. **Gabriel L. Wallau:** Supervision, Conceptualization, Investigation, Writing – review & editing. **Richard S. Salvato**: Supervision, Conceptualization, Investigation, Writing – review & editing.

## DECLARATION OF COMPETING INTEREST

None.

## ACKNOWLEDGMENTS

The authors would like to thank the Secretaria da Saúde do Estado do Rio Grande do Sul for infrastructure availability and the work of professionals in charge of arbovirus control policies and diagnosis.

## DATA AVAILABILITY

Raw reads and consensus fasta sequences from coinfection cases are available in the GISAID database under accession codes: EPI_ISL_18907330, EPI_ISL_18907331, EPI_ISL_18907332, EPI_ISL_18907333, EPI_ISL_18907334, and EPI_ISL_18907335. Datasets used in phylogenetic analysis, major and minor consensus sequences, and genome assembly metrics are available on Github (https://github.com/salvatolab/coinfection-Dengue-clades-Southern-Brazil).

## SUPPLEMENTARY MATERIALS

**Supplementary Figure 1:**
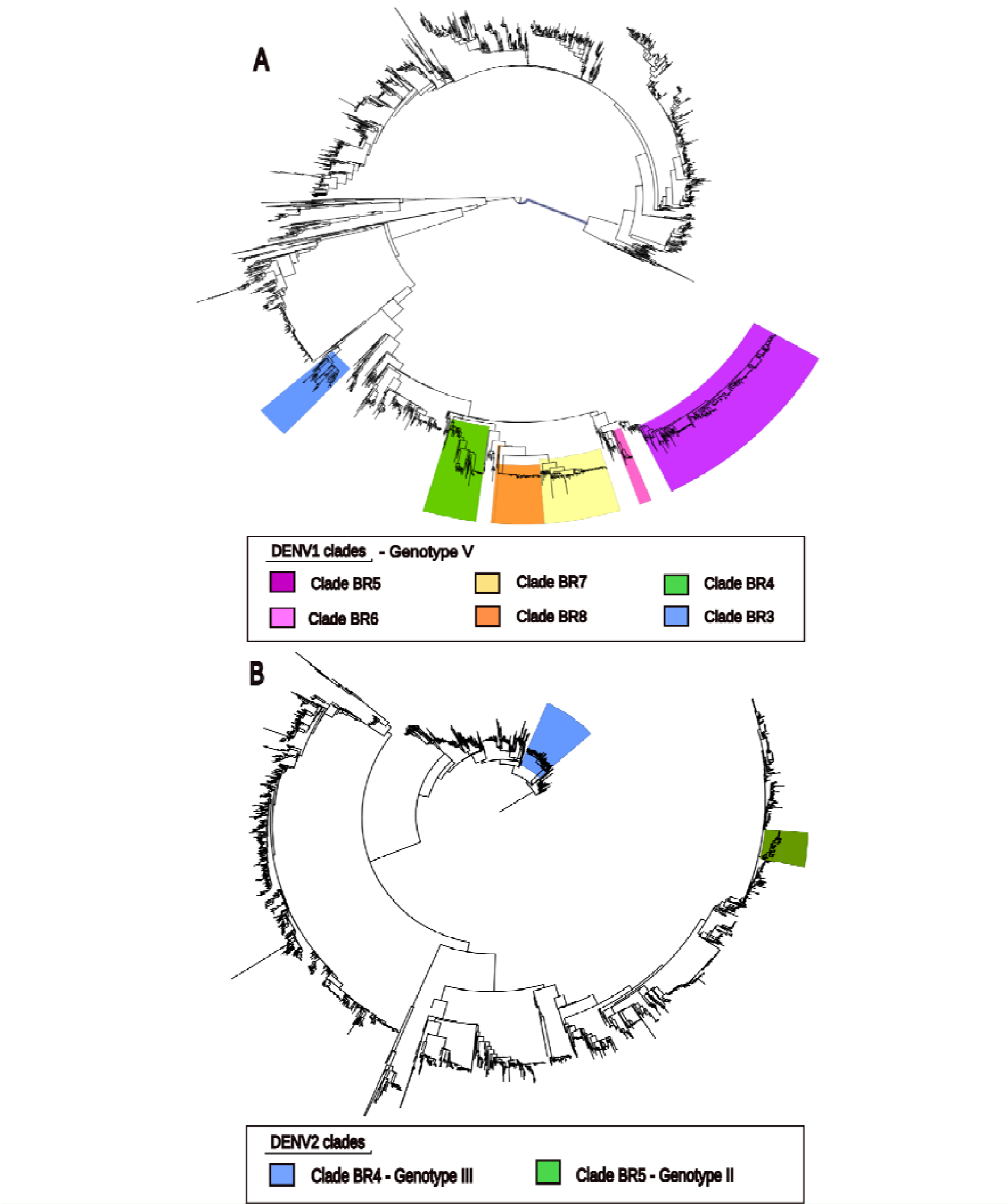
**A**. Maximum likelihood tree for complete dataset DENV-1. The six clades identified in our study are highlighted, as shown in the legend below the tree; **B**. DENV-2 maximum likelihood tree for the complete dataset. The two clades identified in our study are highlighted, as shown in the legend below the tree.

**Supplementary Figure 2:**
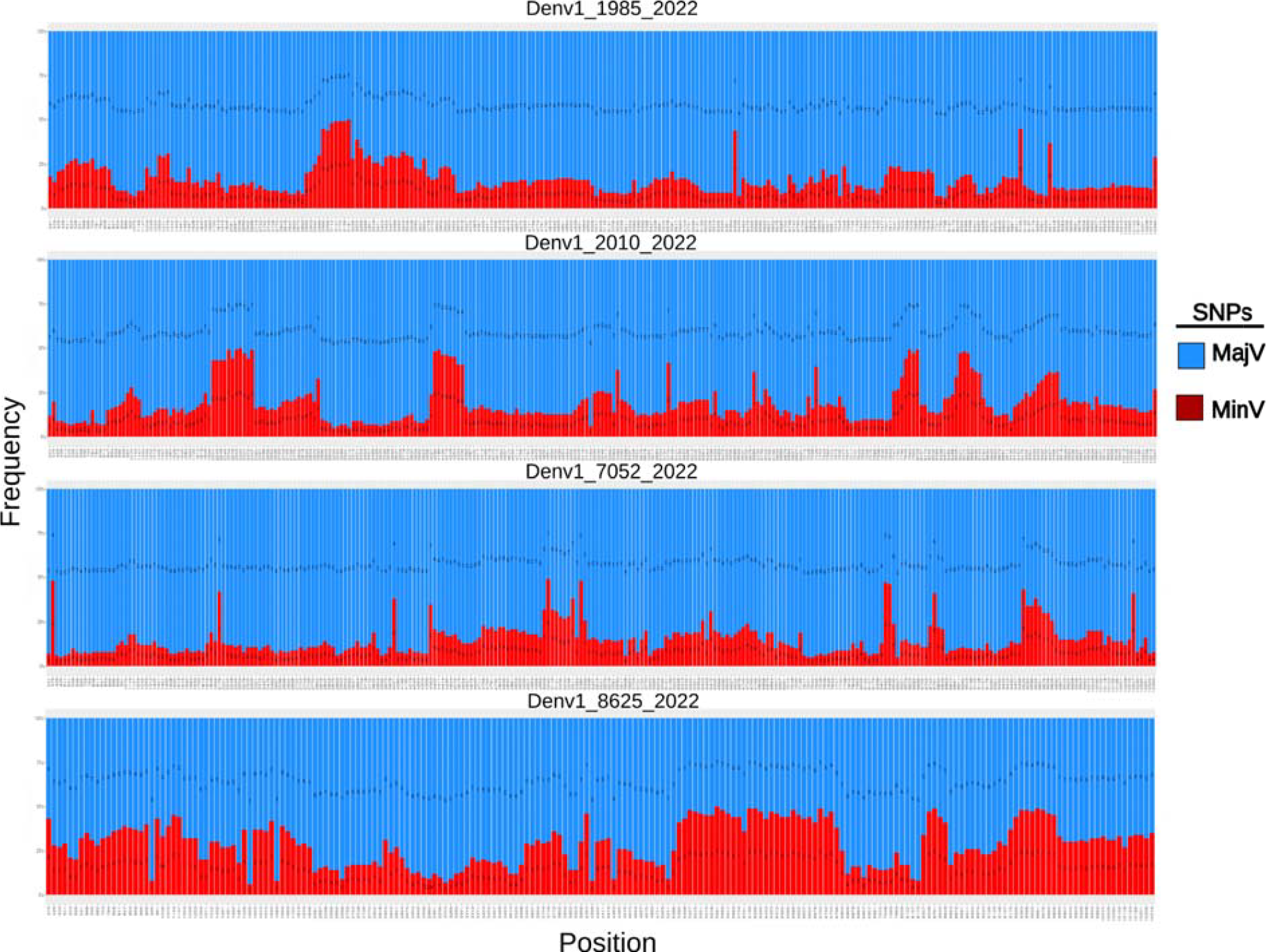
Plot with the frequencies of the two well-supported alleles for the MajV and MinV genomes of coinfection cases: Denv1_1985_2022, Denv1_2010_2022, Denv1_7052_2022 and Denv1_8625_2022. On the X axis, we have the position in the genome; on the Y axis, we have the frequency in percentage over the depth of reads in a given allele. The part of the bar in blue represents the frequency of the MajV allele, and the part in red represents the frequency of the MinV allele in percentage. The nucleotide representing this frequency in each position is positioned in the center of the bar.

**Supplementary Figure 3.**
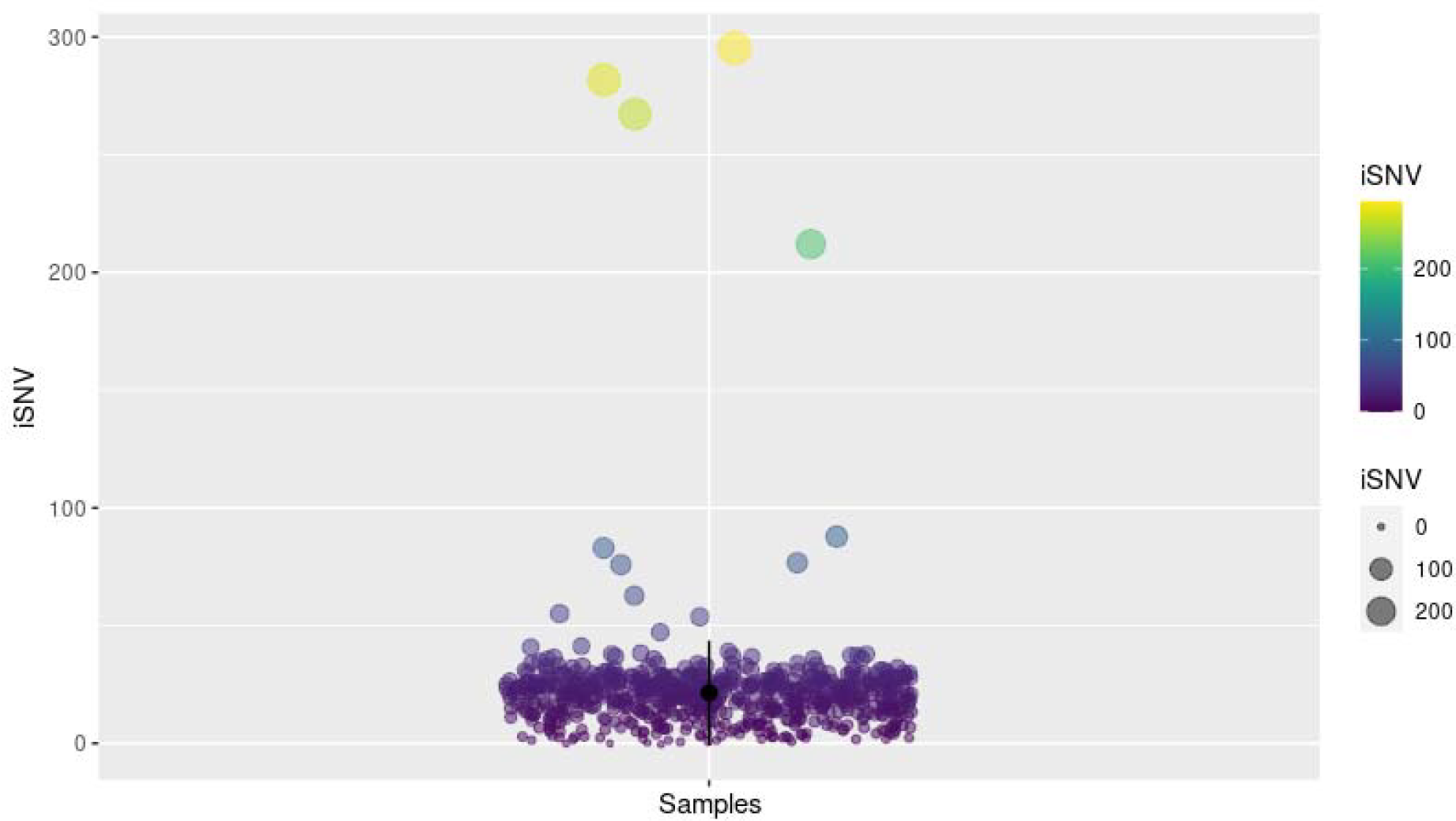
Plot with iSNV values for complete genomes of all samples, highlighting the four outlier samples with a notable gap between the values.

**Supplementary Table 1.** iSNV values for complete genomes of all samples. The first column contains all 627 DENV1 samples followed by their iSNV values and all 40 DENV2 samples followed by their iSNV values.

## Notes

https://github.com/salvatolab/coinfection-Dengue-clades-Southern-Brazil

### Competing Interest Statement

The authors have declared no competing interest.

## REFERENCES

Barreto, M.L., Teixeira, M.G., 2008. Dengue in Brazil: Epidemiological situation and Contribution to a Research Agenda. Estudos avançados.

Brasil. Ministério da Saúde. DATASUS - Doenças e Agravos de Notificação - 2007 em diante (SINAN). Available at: https://datasus.saude.gov.br/acesso-a-informacao/doencas-e-agravos-de-notificacao-de-2007-em-diante-sinan

Brito, A.F., Machado, L.C., Oidtman, R.J., Siconelli, M.J.L., Tran, Q.M., Fauver, J.R., Carvalho, R.D.D.O., Dezordi, F.Z., Pereira, M.R., De Castro-Jorge, L.A., Minto, E.C.M., Passos, L.M .R., Kalinich, C.C., Petrone, M.E., Allen, E., España, G.C., Huang, A.T., Cummings, D.A.T., Baele, G., Franca, R.F.O., Da Fonseca, B.A.L., Perkins, T.A., Wallau, G.L., Grubaugh, N.D., 2021. Lying in wait: the resurgence of dengue virus after the Zika epidemic in Brazil. Nat Commun 12, 2619. 10.1038/s41467-021-22921-7

Brito, C.A.A., Azevedo, F., Cordeiro, M.T., Marques, E.T.A., Franca, R.F.O., 2017. Central and peripheral nervous system involvement caused by Zika and chikungunya coinfection. PLoS Negl Trop Dis 11, e0005583. 10.1371/journal.pntd.0005583

Carrillo-Hernández, M.Y., Ruiz-Saenz, J., Villamizar, L.J., Gómez-Rangel, S.Y., Martínez-Gutierrez, M., 2018. Co-circulation and simultaneous co-infection of dengue, chikungunya, and zika viruses in patients with febrile syndrome at the Colombian-Venezuelan border. BMC Infect Dis 18, 61. 10.1186/s12879-018-2976-1

Codeco, C.T., Oliveira, S.S., Ferreira, D.A.C., Riback, T.I.S., Bastos, L.S., Lana, R.M., Almeida, I.F., Godinho, V.B., Cruz, O.G., Coelho, F.C., 2022. Fast expansion of dengue in Brazil. The Lancet Regional Health - Americas 12, 100274. 10.1016/j.lana.2022.100274

Da Silva Neto, S.R., Tabosa De Oliveira, T., Teixiera, I.V., Medeiros Neto, L., Souza Sampaio, V., Lynn, T., Endo, P.T., 2022. Arboviral disease record data - Dengue and Chikungunya, Brazil, 2013–2020. Sci Data 9, 198. 10.1038/s41597-022-01312-7

Descloux, E., Cao-Lormeau, V.-M., Roche, C., De Lamballerie, X., 2009. Dengue 1 Diversity and Microevolution, French Polynesia 2001–2006: Connection with Epidemiology and Clinics. PLoS Negl Trop Dis 3, e493. 10.1371/journal.pntd.0000493

Dezordi, F.Z., Neto, A.M.D.S., Campos, T.D.L., Jeronimo, P.M.C., Aksenen, C.F., Almeida, S.P., Wallau, G.L., on behalf of the Fiocruz COVID-19 Genomic Surveillance Network, 2022a. ViralFlow: A Versatile Automated Workflow for SARS-CoV-2 Genome Assembly, Lineage Assignment, Mutations and Intrahost Variant Detection. Viruses 14, 217. 10.3390/v14020217

Dezordi, F.Z., Resende, P.C., Naveca, F.G., Do Nascimento, V.A., De Souza, V.C., Dias Paixão, A.C., Appolinario, L., Lopes, R.S., Da Fonseca Mendonça, A.C., Barreto Da Rocha, A.S., Martins Venas, T.M., Pereira, E.C., Paiva, M.H.S., Docena, C., Bezerra, M.F., Machado, L.C., Salvato, R.S., Gregianini, T.S., Martins, L.G., Pereira, F.M., Rovaris, D.B., Fernandes, S.B., Ribeiro-Rodrigues, R., Costa, T.O., Sousa, J.C., Miyajima, F., Delatorre, E., Gräf, T., Bello, G., Siqueira, M.M., Wallau, G.L., 2022b. Unusual SARS-CoV-2 intrahost diversity reveals lineage superinfection. Microbial Genomics 8. 10.1099/mgen.0.000751

Furuya-Kanamori, L., Liang, S., Milinovich, G., Soares Magalhaes, R.J., Clements, A.C.A., Hu, W., Brasil, P., Frentiu, F.D., Dunning, R., Yakob, L., 2016. Co-distribution and co-infection of chikungunya and dengue viruses. BMC Infect Dis 16, 84. 10.1186/s12879-016-1417-2

Gräf, T., Ferreira, C.D.N., De Lima, G.B., De Lima, R.E., Machado, L.C., Campos, T.D.L., Schemberger, M.O., Faoro, H., Paiva, M.H.S., Bezerra, M.F., Nascimento, V., Souza, V., Nascimento, F., Mejía, M., Silva, D., De Oliveira, Y.S., Gonçalves, L., Ramos, T.C.A., De Castro, D.B., Arcanjo, A.R., Dantas, H.A.P., Presibella, M.M., Fernandes, S.B., Gregianini, T.S., Paz E Silva, K.M., Sacchi, C.T., Cruz, A.C.R., Duarte Dos Santos, C.N., Bispo De Filippis, A.M., Bello, G., Wallau, G.L., Salvato, R.S., Naveca, F., 2023. Multiple introductions and country-wide spread of DENV-2 genotype II (Cosmopolitan) in Brazil. Virus Evolution 9, vead059. 10.1093/ve/vead059

Gregianini, T.S., Ranieri, T., Favreto, C., Nunes, Z.M.A., Tumioto Giannini, G.L., Sanberg, N.D., Da Rosa, M.T.M., Da Veiga, A.B.G., 2017. Emerging arboviruses in Rio Grande do Sul, Brazil: Chikungunya and Zika outbreaks, 2014L2016. Reviews in Medical Virology 27, e1943. 10.1002/rmv.1943

Gregianini, T.S., TumiotoLGiannini, G.L., Favreto, C., Plentz, L.C., Ikuta, N., Da Veiga, A.B.G., 2018. Dengue in Rio Grande do Sul, Brazil: 2014 to 2016. Reviews in Medical Virology 28, e1960. 10.1002/rmv.1960

Gularte, J.S., Sacchetto, L., Demoliner, M., Girardi, V., Da Silva, M.S., Filippi, M., Pereira, V.M.D.A.G., Hansen, A.W., Da Silva, L.L., Fleck, J.D., De Almeida, P.R., Nogueira, M.L., Spilki, F.R., 2023. DENV-1 genotype V linked to the 2022 dengue epidemic in Southern Brazil. Journal of Clinical Virology 168, 105599. 10.1016/j.jcv.2023.105599

Halstead, S.B., 2014. Dengue Antibody-Dependent Enhancement: Knowns and Unknowns. Microbiol Spectr 2, 2.6.30. 10.1128/microbiolspec.AID-0022-2014

Kading, R.C., Brault, A.C., Beckham, J.D., 2020. Global Perspectives on Arbovirus Outbreaks: A 2020 Snapshot. TropicalMed 5, 142. 10.3390/tropicalmed5030142

Kalyaanamoorthy, S., Minh, B.Q., Wong, T.K.F., Von Haeseler, A., Jermiin, L.S., 2017. ModelFinder: fast model selection for accurate phylogenetic estimates. Nat Methods 14, 587–589. 10.1038/nmeth.4285

Katoh, K., Rozewicki, J., Yamada, K.D., 2019. MAFFT online service: multiple sequence alignment, interactive sequence choice and visualization. Briefings in Bioinformatics 20, 1160–1166. 10.1093/bib/bbx108

Ko, H.-Y., Li, Y.-T., Chao, D.-Y., Chang, Y.-C., Li, Z.-R.T., Wang, M., Kao, C.-L., Wen, T.-H., Shu, P.-Y., Chang, G.-J.J., King, C.-C., 2018. Inter- and intra-host sequence diversity reveal the emergence of viral variants during an overwintering epidemic caused by dengue virus serotype 2 in southern Taiwan. PLoS Negl Trop Dis 12, e0006827. 10.1371/journal.pntd.0006827

Larsson, A., 2014. AliView: a fast and lightweight alignment viewer and editor for large datasets. Bioinformatics 30, 3276–3278. 10.1093/bioinformatics/btu531

Machado, L.C., De Morais-Sobral, M.C., Campos, T.D.L., Pereira, M.R., De Albuquerque, M.D.F.P.M., Gilbert, C., Franca, R.F.O., Wallau, G.L., 2019. Genome sequencing reveals coinfection by multiple chikungunya virus genotypes in a recent outbreak in Brazil. PLoS Negl Trop Dis 13, e0007332. 10.1371/journal.pntd.0007332

Marinho, P.E.S., Bretas De Oliveira, D., Candiani, T.M.S., Crispim, A.P.C., Alvarenga, P.P.M., Castro, F.C.D.S., Abrahão, J.S., Rios, M., Coimbra, R.S., Kroon, E.G., 2017. Meningitis Associated with Simultaneous Infection by Multiple Dengue Virus Serotypes in Children, Brazil. Emerg. Infect. Dis. 23, 115–118. 10.3201/eid2301.160817

Marklewitz, M., Junglen, S., 2019. Evolutionary and ecological insights into the emergence of arthropod-borne viruses. Acta Tropica 190, 52–58. 10.1016/j.actatropica.2018.10.006

Mayer, S.V., Tesh, R.B., Vasilakis, N., 2017. The emergence of arthropod-borne viral diseases: A global prospective on dengue, chikungunya and zika fevers. Acta Tropica 166, 155–163. 10.1016/j.actatropica.2016.11.020

Minh, B.Q., Schmidt, H.A., Chernomor, O., Schrempf, D., Woodhams, M.D., Von Haeseler, A., Lanfear, R., 2020. IQ-TREE 2: New Models and Efficient Methods for Phylogenetic Inference in the Genomic Era. Molecular Biology and Evolution 37, 1530–1534. 10.1093/molbev/msaa015

Parameswaran, P., Charlebois, P., Tellez, Y., Nunez, A., Ryan, E.M., Malboeuf, C.M., Levin, J.Z., Lennon, N.J., Balmaseda, A., Harris, E., Henn, M.R., 2012. Genome-Wide Patterns of Intrahuman Dengue Virus Diversity Reveal Associations with Viral Phylogenetic Clade and Interhost Diversity. J Virol 86, 8546–8558. 10.1128/JVI.00736-12

Pollett, S., Fauver, J.R., Maljkovic Berry, I., Melendrez, M., Morrison, A., Gillis, L.D., Johansson, M.A., Jarman, R.G., Grubaugh, N.D., 2020. Genomic Epidemiology as a Public Health Tool to Combat Mosquito-Borne Virus Outbreaks. The Journal of Infectious Diseases 221, S308–S318. 10.1093/infdis/jiz302

Pollett, S., Melendrez, M.C., Maljkovic Berry, I., Duchêne, S., Salje, H., Cummings, D.A.T., Jarman, R.G., 2018. Understanding dengue virus evolution to support epidemic surveillance and counter-measure development. Infection, Genetics and Evolution 62, 279–295. 10.1016/j.meegid.2018.04.032

Rio Grande do Sul. Secretaria Estadual da Saúde. Painel de Casos de Dengue RS. Available at: https://dengue.saude.rs.gov.br

Rodriguez-Morales, A.J., Villamil-Gómez, W.E., Franco-Paredes, C., 2016. The arboviral burden of disease caused by co-circulation and co-infection of dengue, chikungunya and Zika in the Americas. Travel Medicine and Infectious Disease 14, 177–179. 10.1016/j.tmaid.2016.05.004

Romano, C.M., Lauck, M., Salvador, F.S., Lima, C.R., Villas-Boas, L.S., Araújo, E.S.A., Levi, J.E., Pannuti, C.S., O’Connor, D., Kallas, E.G., 2013. Inter- and Intra-Host Viral Diversity in a Large Seasonal DENV2 Outbreak. PLoS ONE 8, e70318. 10.1371/journal.pone.0070318

Shukla, R., Ramasamy, V., Shanmugam, R.K., Ahuja, R., Khanna, N., 2020. Antibody-Dependent Enhancement: A Challenge for Developing a Safe Dengue Vaccine. Front. Cell. Infect. Microbiol. 10, 572681. 10.3389/fcimb.2020.572681

Siqueira, J.B., Martelli, C.M.T., Coelho, G.E., Simplício, A.C.D.R., Hatch, D.L., 2005. Dengue and Dengue Hemorrhagic Fever, Brazil, 1981–2002. Emerg. Infect. Dis. 11, 48–53. 10.3201/eid1101.031091

Thai, K.T.D., Henn, M.R., Zody, M.C., Tricou, V., Nguyet, N.M., Charlebois, P., Lennon, N.J., Green, L., De Vries, P.J., Hien, T.T., Farrar, J., Van Doorn, H.R., De Jong, M.D., Birren, B.W., Holmes, E.C., Simmons, C.P., 2012. High-Resolution Analysis of Intrahost Genetic Diversity in Dengue Virus Serotype 1 Infection Identifies Mixed Infections. J Virol 86, 835–843. 10.1128/JVI.05985-11

Tumioto, G.L., Gregianini, T.S., Dambros, B.P., Cestari, B.C., Alves Nunes, Z.M., Veiga, A.B.G., 2014. Laboratory Surveillance of Dengue in Rio Grande do Sul, Brazil, from 2007 to 2013. PLoS ONE 9, e104394. 10.1371/journal.pone.0104394

Villabona-Arenas, C.J., De Oliveira, J.L., Capra, C.D.S., Balarini, K., Loureiro, M., Fonseca, C.R.T.P., Passos, S.D., Zanotto, P.M.D.A., 2014. Detection Of Four Dengue Serotypes Suggests Rise In Hyperendemicity In Urban Centers Of Brazil. PLoS Negl Trop Dis 8, e2620. 10.1371/journal.pntd.0002620

Villamil-Gómez, W.E., Rodríguez-Morales, A.J., Uribe-García, A.M., González-Arismendy, E., Castellanos, J.E., Calvo, E.P., Álvarez-Mon, M., Musso, D., 2016. Zika, dengue, and chikungunya co-infection in a pregnant woman from Colombia. International Journal of Infectious Diseases 51, 135–138. 10.1016/j.ijid.2016.07.017

Wang, W.-K., Lin, S.-R., Lee, C.-M., King, C.-C., Chang, S.-C., 2002. Dengue Type 3 Virus in Plasma Is a Population of Closely Related Genomes: Quasispecies. J Virol 76, 4662–4665. 10.1128/JVI.76.9.4662-4665.2002

